# DECIPHERING THE DARK SIDE OF HISTONE ADP-RIBOSYLATION: WHAT STRUCTURAL FEATURES OF DAMAGED NUCLEOSOME REGULATE THE ACTIVITIES OF PARP1 AND PARP2

**DOI:** 10.1101/2025.01.29.635606

**Authors:** Kurgina Tatyana A., Moor Nina A., Kutuzov Mikhail M., Endutkin Anton V., Lavrik Olga I.

## Abstract

Poly(ADP-ribose) polymerases are critical enzymes contributing to regulation of numerous cellular processes, including DNA repair. Within the PARP family, PARP1 and PARP2 primarily facilitate PARylation in the nucleus, particularly responding to genotoxic stress. The activity of PARPs is influenced by the nature of DNA damage and multiple protein partners, with HPF1 being the important one. Forming a joint active site with PARP1 (PARP2), HPF1 contributes to histone PARylation and following chromatin remodelling during genotoxic stress events. This study elucidates interrelation between the presence and location of a one-nucleotide gap within the nucleosome core particle (NCP) and PARP activities in automodification and heteromodification of histones. Utilizing a combination of classical biochemical methods with fluorescence-based technique and a single-molecule mass photometry approach, we have shown that the NCP architecture impacts the efficiency and pattern of histone ADP-ribosylation and binding to the histones-associated damaged DNA more significantly for PARP2 than for PARP1. Analysis based on existing studies of HPF1-dependent ADP-ribosylome and NCP structural dynamics allows to suggest that the DNA damage location and the conformational flexibility of histone tails modulated by post-translational modifications are crucial for delineating the distinct roles of PARP1 and PARP2 during genotoxic stress responses.

**GRAPHICAL ABSTRACT:** 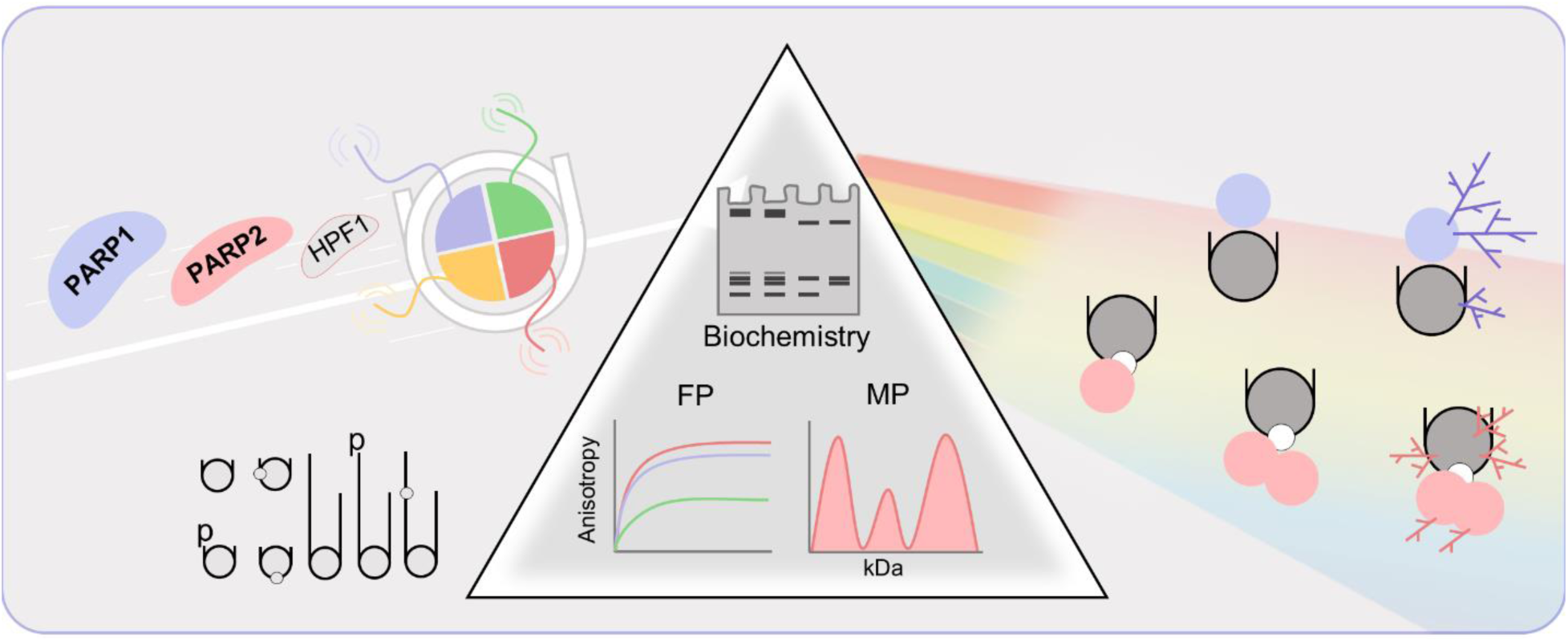

## INTRODUCTION

Poly(ADP-ribosyl)ation (PARylation) is a dynamic post-translational modification of biomolecules such as proteins and nucleic acids, catalysed by members of the PARP family. PARPs possess a conserved catalytic domain that facilitates the binding of nicotinamide adenine dinucleotide (NAD^+^) and the transfer of ADP-ribose moieties from the donor NAD^+^ to various amino acid residues on protein targets. PARPs can modify themselves (automodification) or other target molecules (heteromodification) (1).

Among the PARP family members, PARP1 and PARP2 are responsible for PAR synthesis in the nucleus and play pivotal roles in regulating multiple cellular processes, including DNA repair, replication, and modulation of chromatin architecture (2−6). Covalent PARylation of proteins can impact their function, localization, and stability. Furthermore, the highly negatively charged PAR polymer can serve as a molecular scaffold, recruiting and concentrating various proteins, such as DNA repair factors, chromatin remodelers, and transcription factors (5,7−10). Thus, PARPs can regulate functions of proteins both through the covalent PARylation and facilitation of their co-localization at damaged DNA.

Despite extensive research on PARylation, many aspects of this process remain unclear. The initiation of PARylation involves the coordination of NAD^+^ and the acceptor amino acid of the target protein at the enzyme’s active site. The complexity of this coordination likely leads to hydrolysis of NAD^+^ and synthesis of free PAR chains before the successful initiation of covalent PARylation (11−15). The efficiency of initiation depends on several factors, including the interaction of the acceptor protein with PARP, the flexibility of the modified region, and the nature of the target amino acid residues and their surrounding environment. Many studies have focused on the role of PAR- and DNA-mediated interactions or direct protein-protein interactions between PARP and its target proteins. The major mechanism of selective PARylation is constituted by PAR-binding domains in target proteins (16−18), but the interaction mediated by a specific DNA structure may also contribute to the substrate targeting (19). However, the factors influencing specific selection of acceptor amino acids are not fully understood. It is believed that while a consensus sequence for the selection of PARylation sites may not be required, certain preferences exist. For instance, ADP-ribosylation of glutamate follows a proline-directed motif (PXE*, EP, PXXE) (20), while serine ADP-ribosylation, catalysed by PARP1/PARP2 in complex with histone PARylation factor 1 (HPF1), preferentially targets the KS motif (21,22). Thus, both the steric accessibility of PARylation sites and the primary structure of the target protein influence trans-PARylation, though the precise mechanisms remain to be clarified.

Core histones (H2A, H2B, H3, and H4) are well-established substrates for HPF1-dependent ADP-ribosylation (23−25). Proteomic studies demonstrate that histones are among the most abundant acceptors of Ser-linked ADP-ribosylation upon oxidative stress (26). This is likely associated with HPF1-dependent PARylation of histones at sites of DNA damage, which promotes chromatin relaxation (4,27). In this context, it is important to highlight the distinct roles of PARP1 and PARP2. *In vivo* experiments with PARP1 depletion show that, in the absence of PARP1, histone PARylation and chromatin relaxation are significantly reduced (27,28). In contrast, PARP2 knockout does not produce a similar effect (28,29). Thus, the predominant role of PARP1 in initiating the DNA damage response is clear, while the role of PARP2 appears less significant. On the other hand, PARP1 and its catalytic activities facilitate the enrichment of PARP2 at DNA damage foci (30,31). Therefore, the reduction in histone PARylation in the absence of PARP1 may, among other factors, be linked to impaired recruitment of PARP2 to DNA damage sites. In other words, there is a potential gap in our understanding of the specific role of PARP2 in the genotoxic stress response. Interestingly, *in vitro* studies indicate that PARP2 modifies histones more efficiently than PARP1 and is specifically activated by BER-related damage in the nucleosome core particle (NCP) (13,32). In the current study, we further compare the roles of PARP1 and PARP2 in HPF1-dependent histone modification.

The structured histone domains serve as a foundation for the assembly of stable NCP, while the flexible histone tails at N- and C-termini play a crucial role in regulating interactions between the nucleosome and various factors (25). Furthermore, the histone tails are critically involved in the modulation of histone-DNA interactions (i.e. nucleosome dynamics) via multiple post-translational modifications (9,33−36). Consequently, the well-established sites of various chemical modifications and ADP-ribosylation are predominantly found within the histone tails.

This research specifically investigates how the architecture of the PARP-NCP complex influences the HPF1-dependent PARylation of histones. The use of NCP assembled *in vitro* on the Widom-603 DNA provides an advantageous model for studying heteroPARylation. The known dynamics of histone tails within the NCP, coupled with the precise positioning of DNA on the histone core facilitated by the Widom-603 sequence, offer clarity in the experimental setup. Furthermore, the position of a DNA damage (one-nucleotide gap intermediate of base excision repair, BER) shown in our previous research to influence activities of PARP1/PARP2 in the auto- and heteromodification reactions (32) can be precisely determined in the compact NCP structure.

The present results highlight dependence of the extent and pattern of histone ADP-ribosylation on the presence of BER-specific DNA damage in distinct superhelical locations (SHLs) or outside the NCP. PARP1 and PARP2 revealed different sensitivity to the structural features of NCP, further suggesting their specific roles in DNA damage response and chromatin remodelling.

## MATERIALS AND METHODS

### Materials

Core histones were isolated from *Gallus gallus* erythrocytes and purified as described previously (37). Recombinant wild-type human APE1, human PARP1, murine PARP2 and human HPF1 were expressed and purified as detailed previously (13,38,39). *E. coli* uracil-DNA glycosylase (UDG) was from Biosan (Novosibirsk, Russia). The recombinant bovine poly(ADP-ribose) glycohydrolase (PARG) generously provided by E. Ilina (ICBFM, Novosibirsk, Russia) was purified as described previously (40). Recombinant polymerase RB69 and *S. pyogenes* Cas9 kindly provided by D. Zharkov (ICBFM, Novosibirsk, Russia) were purified as described previously (41,42). DNA primers for synthesis of Widom603 147 bp DNA were synthesized by Lumiprobe (Moscow, Russia). pGEM-3z/603 (Addgene plasmid #26658; http://n2t.net/addgene:26658; RRID: Addgene_26658) was a gift from J. Widom. pBR322 (Addgene plasmid #1979) was from SibEnzyme (Novosibirsk, Russia). The ^32^P-labelled NAD^+^ was synthesized enzymatically following a described method (43), using [α-^32^P]ATP (with specific activity of 3000 Ci/mmol, synthesized by Laboratory of Biotechnology (ICBFM, Novosibirsk, Russia). NAD^+^, reagents for electrophoresis and basic components of buffers were purchased from Sigma-Aldrich (USA).

### Methods

#### Preparation of DNA and Nucleosome

The amplification of Widom603 147 bp DNA and subsequent reconstitution of nucleosomes using the Widom603 sequence were performed as described previously (37,44). Briefly, DNA was obtained by PCR using different primers (Table 1); nucleosome was reconstituted by dialysis of DNA-histones mixture against the NaCl concentration gradient from 2 M to 10 mM. The homogeneity of the nucleosome sample was analyzed by the electrophoretic mobility shift assay on a 4% nondenaturing PAG. DNA (both free and nucleosome-associated) containing a one-nucleotide gap (gap-DNA, gap-NCP) was prepared and characterized as described previously (32).

**Table 1.**
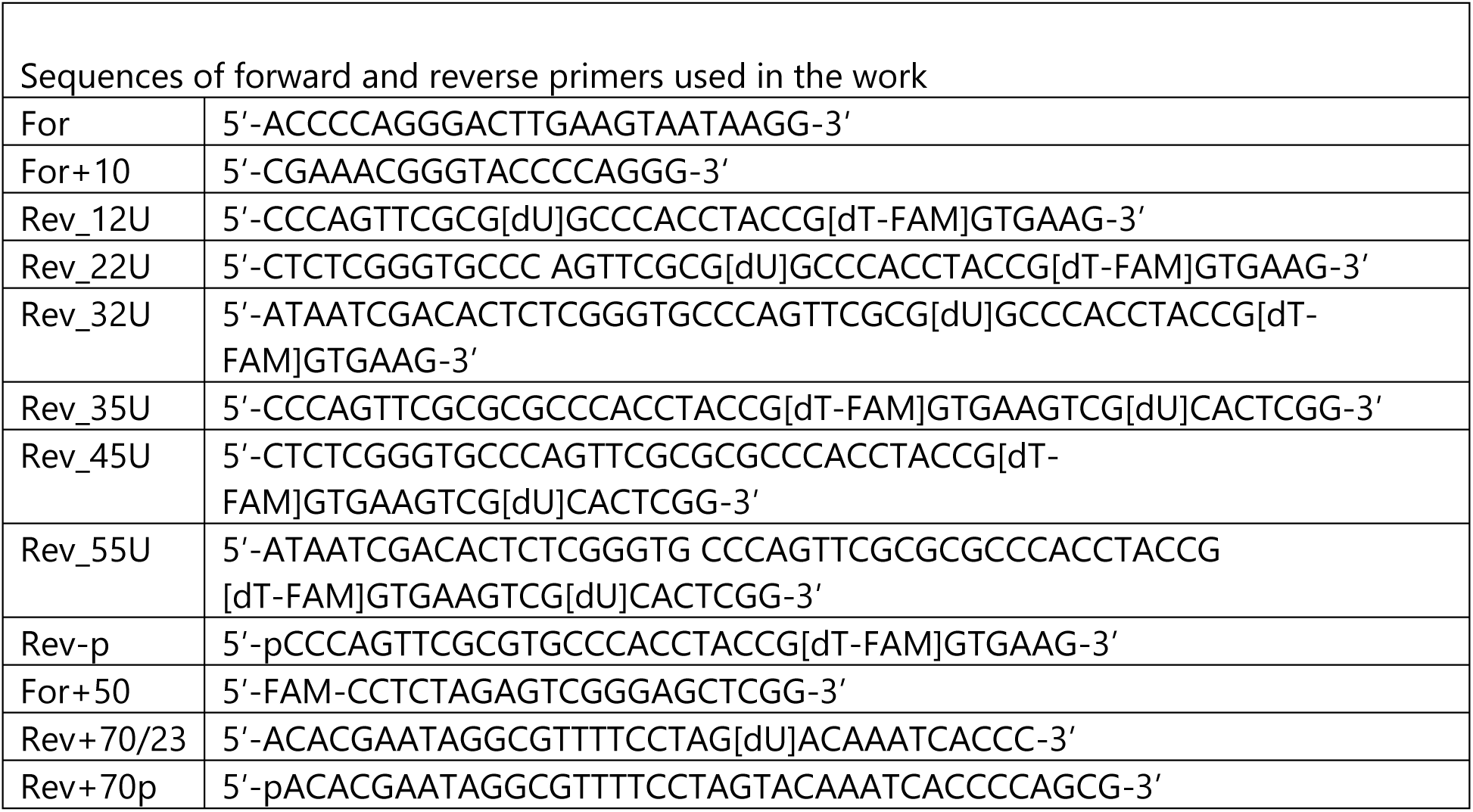
Sequences of forward and reverse primers used in the work

#### Testing of PARP Activity in the Poly(ADP-ribose) Synthesis

Catalysed by PARP1 and PARP2 autopoly(ADP-ribosyl)ation and covalent labelling of histones were carried out in a standard 10 μl reaction mixture containing 50 mM Tris-HCl, pH 8.0, 50 mM NaCl, 5 mM MgCl_2_, 1 µM [^32^P]NAD^+^, 250 nM DNA/NCP (or gap-DNA/gap-NCP), 500 nM PARP1 (PARP2), and 1 μM HPF1. The reaction was initiated by adding [^32^P]NAD^+^ to a protein-DNA mixture preassembled on ice. After incubating the mixtures at 37°C for 45 min, the reactions were terminated by the addition of SDS-PAGE sample buffer and heating for 3 min at 95°C. To perform PAR hydrolysis, the ADP-ribosylation reaction was stopped by addition of 1 μM olaparib and 10 mM EDTA, and the mixture was further incubated with 1 μM PARG for 1 h at 37°C before addition of SDS-PAGE sample buffer. The reaction products were separated by 20% SDS-PAGE (a ratio between acrylamide and bis-acrylamide of 99:1); bands of proteins labelled with [^32^P]ADP-ribose were analyzed by using the Typhoon imaging system (GE Healthcare Life Sciences) and Quantity One Basic Software (Bio-Rad). The radiolabelled signals of modified proteins were quantified as follows: the total (raw) signal of the smeared band of modified protein (indicated for each protein in the autoradiograms) was quantified and the same-size background signal of gel in the respective lane was subtracted from the raw signal. The quantitative data presented in histograms were obtained in at least three independent experiments.

#### Fluorescence Studies of PARP1/PARP2 Interaction with DNAs/NCPs

In direct titration experiments, fluorescence anisotropy measurements of labelled DNAs (free or nucleosome-associated) were performed in the absence and presence of various concentrations of the potential protein partner (45). A mixture containing 3 nM 25-FAM-labelled DNA (NCP) and 1–60 nM PARP1 or PARP2 in a binding buffer (50 mM NaCl, 50 mM Tris–HCl, pH 8.0, 5 mM MgCl_2_ and 5 mM DTT) was prepared on ice in a 384-well plate and incubated at room temperature for 10 min. In competition binding experiments, reaction mixtures contained 50 mM NaCl, 50 mM Tris-HCl (pH 8.0), 5 mM DTT, 5 mM MgCl_2_, 10 nM DNA, 5 nM PARP1, and increasing concentrations of plasmid DNA pBR322. The fluorescent probes were excited at 482 nm (482–16 filter plus dichroic filter LP504), and the fluorescence intensities were detected at 530 nm (530–40 filter) to measure the fluorescence anisotropy of FAM. Each measurement consisted of 50 flashes per well, and the resulting values of fluorescence anisotropy were automatically averaged. The measurements in each well were done 3 times with intervals of 1 min.

The average values were used for the final plot, and EC_50_ values were determined with the MARS Data Analysis software (BMG LABTECH). The data were plotted (F vs C) and fitted by four-parameter logistic equation: F = F_0_ + (F_∞_ - F_0_)/[1 + (EC_50_/C)^n^], where F is the measured fluorescence anisotropy of a solution containing the labelled DNA at a given concentration (C) of PARP1/PARP2, F_0_ is the fluorescence anisotropy of solution of the labelled DNA alone, F_∞_ is the fluorescence anisotropy of the labelled DNA saturated with the protein, EC_50_ is the protein concentration at which F - F_0_ = (F_∞_ - F_0_)/2, and n is the Hill coefficient, which denotes the slope of the nonlinear curve. CC_50_ values were determined by fitting average F values by four-parameter logistic equation: F = F_0_ + (F_∞_ - F_0_)/[1 + (CC_50_/C)], where F is the measured fluorescence anisotropy of a solution containing the preformed complex of labelled DNA with PARP1/PARP2 at a given concentration (C) of pBR322, F_0_ is the fluorescence anisotropy of the labelled DNA•PARP1/PARP2 complex alone, F_∞_ is the fluorescence anisotropy of the free labelled DNA, CC_50_ is the concentration of the competitive plasmid pBR322 at which F - F_0_ = (F_∞_ - F_0_)/2. Binding curves presented below and in Extended Data show the best fits of the respective equation, with *R^2^* values matching or exceeding 0.97.

#### Mass Photometry Measurements of PARP1/PARP2 Interaction with NCPs

Formation of PARP1 and PARP2 complexes with NCP147 and gap35-NCP147 was carried out in a 20 μl reaction mixture containing 50 mM Tris-HCl, pH 8.0, 50 mM NaCl, 5 mM MgCl_2_, 3 nM nucleosome (NCP147 or gap35-NCP147) and 6 nM PARP1 (PARP2). The buffer was preliminary ultrafiltered by using Sartorius vivaspin columns with 1,000 kDa pore size PES membrane filter. The reaction mixture was prepared on ice, centrifuged for 10 min at 10,000 rpm (4°C), and incubated at room temperature for 10 min. MP measurements were performed using a OneMP mass photometer (Refeyn, UK). Following the standard protocol (46), 18 μL buffer was loaded to the sample chamber and the objective was focused by using autofocus function, then 2 μL sample was added to the chamber and mixed by pipetting. Data were collected for one minute using the AcquireMP software (Refeyn, UK). All samples were measured using the expanded detection area. The MP signals were calibrated using BSA (69 kDa) (Sigma-Aldrich, USA), recombinant RB69 protein (107 kDa) and recombinant Cas9 protein (158 kDa). MP data were processed with the DiscoverMP software (Refeyn, UK) to calculate relative molecular species populations from the areas of the Gaussian peaks (representing free NCP and its complexes with one or two PARP molecules). The number of counts for each species was estimated from 3 independent experiments to obtain the average species concentration fractions (Extended Data). These data were further processed with program “Kd calculation” as described previously (46,47).

## RESULTS

### The DNA damage location in the nucleosome core particle affects HPF1-dependent ADP-ribosylation of histones

The efficiency of PARP1 and PARP2 catalysed histone PARylation was revealed in our previous study to depend on the presence of an incised AP-site in the nucleosomal DNA (32). This prompted us to further explore possible influence of location of the BER specific DNA damage relative to the DNA blunt ends and the nucleosome core histones on the activities of these enzymes in auto- and heteromodification reactions. The sequence of DNA clone 603, comprising 147 bp, provides precise positioning of DNA within the nucleosome, enabling to study the impact of the DNA damage position (44). The interaction of PARP1 and PARP2 enzymes with nucleosome core particle (NCP) was previously shown being sensitive to the DNA damage orientation (inwards or outwards) within NCP (48). In this study, we constructed NCPs containing uracil (which can be removed enzymatically to create the one-nucleotide gap) at positions 12 and 35 relative to the 603 sequence boundary (SHL 3.5 and 5.5, respectively) to ensure the outwards orientation. Seven NCP variants used in the experiments (Fig. 1a) differ from each other by the presence of 5’-phosphate, one-nucleotide gap (the incised AP site) at the two different positions, and 50/70 bp linkers without or with one-nucleotide gap.

**Fig. 1.**
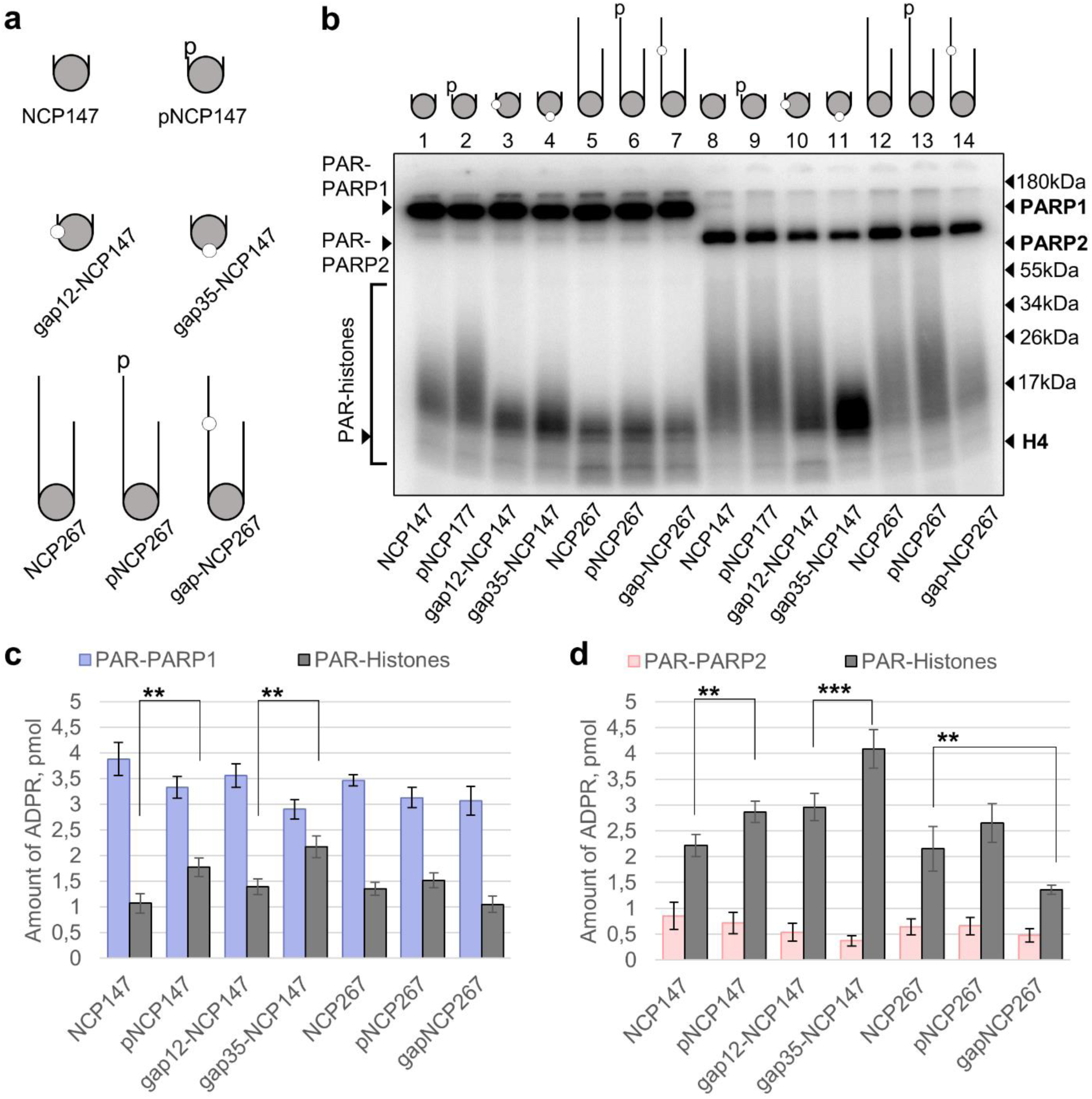
HPF1-dependent ADP-ribosylation of histones is influenced by the position of a BER specific DNA damage: **a** – Schemes of nucleosome structures used in the work; **b –** Autoradiograms show covalent binding of ^32^P-labelled ADP-ribose to PARP1/PARP2 and histones after incubation of each PARP (500 nM) with [^32^P]NAD^+^ (1 μM), HPF1 (1 μM) and NCP with a defined structure (250 nM). Positions of ADP-ribosylated proteins and their native forms (and molecular weight markers) in 20% SDS-PAG are indicated on the left and right of the autoradiograms; **c, d** − Histograms show the amount of ADP-ribose attached to PARP1/PARP2 and histones in the distinct samples. Statistically significant differences in the levels of histone PARylation in the presence of different NCPs (gap12-NCP147 vs gap35-NCP147; NCP267 vs gap-NCP267) are marked: *p* < 0.01 (**), *p* < 0.001 (***).

PARP1 and PARP2 were activated by incubation with different NCP variants in the presence of [^32^P]NAD^+^ and HPF1, and the reaction products were separated by SDS-PAGE to compare the levels of automodification and heteromodification of histones with ADP-ribose (ADPR) (Fig. 1b). We have to emphasize that low concentrations of NAD^+^ were utilized in order to prevent the formation of long PAR chains, which can significantly change the electrophoretic mobility of modified proteins. The primary target for modification catalysed by PARP1 was the enzyme itself, irrespective of the NCP structure used. The amount of ADPR bound to PARP1 in all samples was 1.3-4-fold higher compared to that bound to histones (Fig. 1c). An increased level of histone modification was detected when the nucleosomal DNA contained 5’-phosphate or one-nucleotide gap (pNCP147, gap12-NCP147, gap35-NCP147 vs NCP147), with the highest yield of heteromodification reaction being observed in the presence of NCP bearing the gap at position 35. The addition of 50 bp and 70 bp linkers (without or with 5’-phosphate or gap) to NCP147 (NCP267, pNCP267, gapNCP267) produced insignificant effects on the total levels of auto- and heteromodification, but noticeably decreased the length of PAR attached to histones (Fig. 1b, lanes 5−7). The data suggest that the presence of linkers modulates the interaction mode of PARP1 with NCP (due to an increased distance between the DNA end and nucleosome core or changes in the entry-exit region of NCP), which in turn influences the enzyme activity in heteromodification reaction at the elongation step.

In contrast to PARP1, PARP2 was more active in modification of histones than in automodification as evidenced by 2.5−10-fold higher yields of the heteromodification reaction (Fig. 1d). The ratio between the levels of auto- and heteromodification reactions as well as the length of PAR attached to histones were highly dependent on the NCP structure used for PARP2 activation. The presence of 5’-phosphate or one-nucleotide gap in the nucleosomal DNA favored the modification of histones; the most efficient histone modification was detected when NCP utilized for PARP2 activation contained the gap at position 35. Thus, PARP2-catalysed modification of histones depends on the position of DNA damage (incised AP site). No significant impact of 50 bp and 70 bp linkers on the relative levels of automodification and histone modification was detected (compare data for NCP147 vs NCP267 and pNCP147 vs pNCP267). However, the introduction of gap into the DNA linker, 47 base pairs away from the histone core, significantly reduced the level of histone modification. Such an effect can result from primary binding of PARP2 outside the nucleosome core due its high affinity for the gap.

### Core Histones in Close Proximity to the DNA Damage are Targets of PARP2-Catalysed Modification

As we have found, the efficiency of PARP1/PARP2 catalysed histone modification depends on the location of one-nucleotide gap in the NCP structure. It was interesting to clarify whether this dependence is related to selective modification of distinct core histones. The HPF1-dependent PARylation reaction catalysed by PARP1/PARP2 was performed in the presence of gap12-NCP147 or gap35-NCP147, which revealed impact of the DNA damage position on the total level of histone modification catalysed by both PARP1 and PARP2. Then ADP-ribose oligomers and polymers were degraded with PARG treatment to produce mono-ADP-ribosylated proteins. Mono-ADP-ribosylation does not change the electrophoretic mobility of proteins in SDS-PAG, enabling their identification (Fig. 2a).

**Fig. 2.**
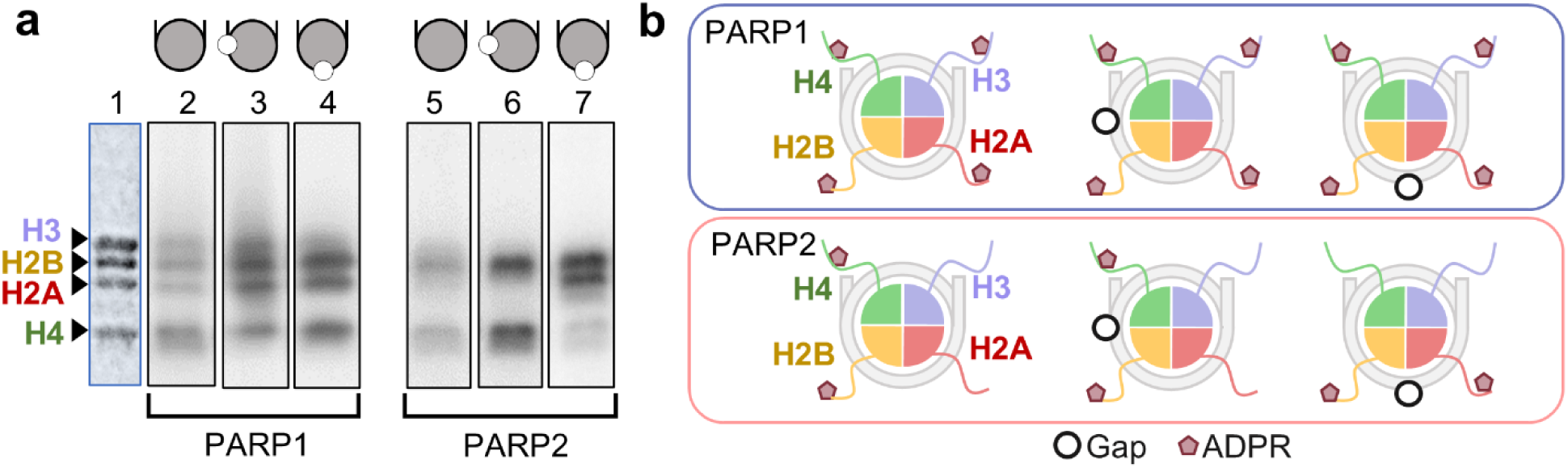
Dependence of histone ADP-ribosylation pattern on the NCP structure: **a** − Autoradiograms show covalent binding of ^32^P-labelled ADP-ribose to histones after incubation of PARP1/PARP2 (500 nM) with [^32^P]NAD^+^ (1 μM), HPF1 (1 μM) and NCP, gap12-NCP or gap35-NCP (250 nM) with following PARG-catalysed hydrolysis and separation of products in 20% SDS-PAG. Gel positions of coomassie-stained histones superposed to their labelled modification products are indicated on the left of the autoradiogram; **b** – Schematic presentation of ADP-ribose attachment to distinct core histones within different NCP structures.

The data obtained indicate that PARP1 catalyzes modification of all four core histones independently on the NCP structure; the main effect of the gap presence in DNA is noticeable increase in the number of initiation events upon the heteroPARylation reaction (Fig. 2a, lines 2−4). In the case of PARP2 no modification of H3 histone was detected (Fig. 2a, lines 5−7). The presence of gap increased the number of PARylation initiation events, with the position of gap having effect on the histone modification pattern. In gap12-NCP structure as well as in NCP, H2B and H4 histones were main acceptors of ADPR, while H2A and H2B histones were primarily modified in gap35-NCP structure. Importantly, known ADP-ribosylation sites on histones H2A and H2B tails (Ser2 and Ser6/Ser14, respectively) are localized close to nucleotide 35 (SHL 3.5), while histones H3 and H4 tails comprising known ADP-ribosylation sites (Ser10/Ser28 and Ser2, respectively) are localized nearer to nucleotide 12 (SHL 5.5) (25,49). Combined, our results suggest that selectivity of HPF1-dependent histone ADP-ribosylation catalysed by PARP2 is provided primarily by the high-affinity enzyme interaction with the one-nucleotide gap and its location relative to the nucleosome core.

To further examine whether the distance between the DNA end and the gap position may impact the efficiency of histone ADP-ribosylation, we designed additional structural variants of gap12-NCP and gap35-NCP containing 10 bp or 20 bp linkers (Fig. 3). The location of gap relative to the core histones remains fixed in two series of variants (gap12-NCP147, gap12-NCP167, gap12-NCP177 and gap35-NCP147, gap35-NCP167, gap35-NCP177), while the distance between the DNA blunt ends and gap varies. Control experiments with length variants of non-gapped NCP (NCP147, NCP167, NCP177) revealed no effects of linkers on the yields of auto- and heteromodification reactions catalysed by PARP1/PARP2 (Fig. 3a,b, lanes 1−3). Similar data were obtained when the length variants of gap12-NCP or gap35-NCP were compared. The levels of histone modification catalysed by both PARP1 and PARP2 in the presence of all three variants of gap35-NCP were nearly equal to each other but higher compared to those in the presence of gap12-NCP variants (Fig. 3c,d). Evidently, the efficiency of histone ADP-ribosylation depends first of all on the DNA gap location relative to the histone core.

**Fig. 3.**
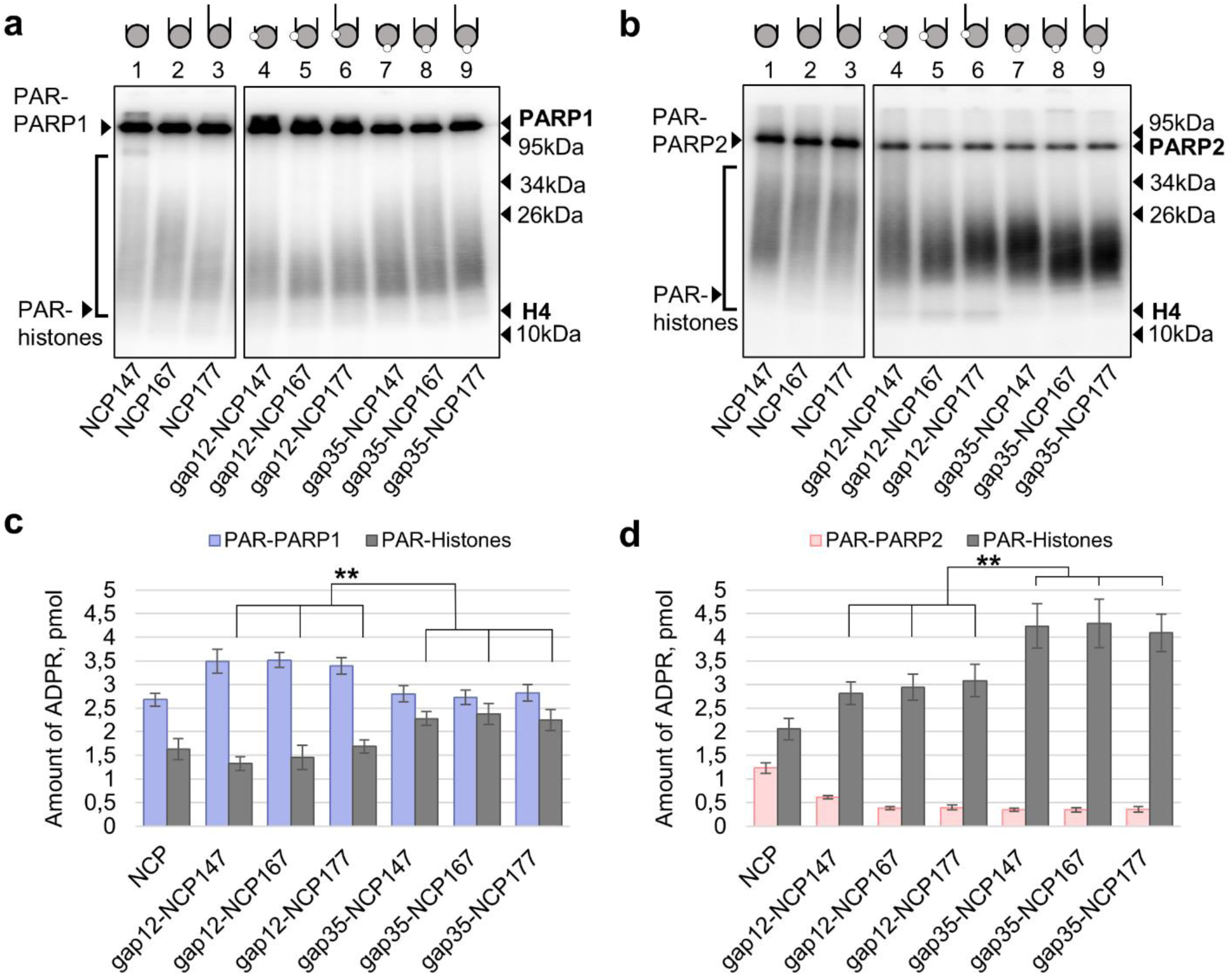
The location of BER-specific DNA damage within NCP determines the efficiency of histone ADP-ribosylation. **a**, **b** – Autoradiograms show covalent binding of ^32^P-labelled ADP-ribose to PARP1, PARP2 and histones after incubation of PARP1/PARP2 (500 nM) with [^32^P]NAD^+^ (1 μM), HPF1 (1 μM) and NCP (250 nM) specified for each sample, and further separation of products in 20% SDS-PAG. Gel positions of PARylated proteins and their native forms (and molecular weight markers) are indicated on the left and right of the autoradiograms; **c, d** − Histograms show the amount of ADP-ribose attached to PARP1/PARP2 and histones in the distinct samples. The total amount of PAR and its distribution between PARP1/PARP2 and histones is shown. Statistically significant differences in the levels of histone PARylation in the presence of three variants of gap12-NCP compared to those of gap35-NCP are marked: *p* < 0.01 (**)

### Impact of DNA Structure on the Interaction of PARP1 and PARP2 with NCP

The dependence of histone ADP-ribosylation on the location of DNA damage raises question whether it is related to different affinities of PARP1/PARP2 for the NCP structural variants. The apparent equilibrium dissociation constants of these complexes were determined using fluorescence anisotropy measurements. The respective DNAs containing one-nucleotide gap at position 12 or 35 and their variants with linkers (Fig. 1a) were labelled with FAM fluorophore at position 25. Taking into account binding of PARP1/PARP2 to undamaged DNA with a lower affinity than to DNA lesions (50,51), the titration experiments were performed in restricted protein concentration range (up to 50−60 nM). In these conditions, contribution of PARP interaction with undamaged DNA to the apparent dissociation constant (EC_50_ value) may be diminished. The EC_50_ values determined for PARP1/PARP2 complexes with different ligands are summarized in Table 2. Their analysis shows that PARP1 binds all gapped DNA variants with very similar affinities and 1.8-fold stronger (*p*˂0.02) than the nongapped DNA, suggesting contribution of gap as a binding site to the affinity. However, we revealed no statistically significant effect of gap presence on the strength of PARP1 binding to NCP ligands, that may result from a more compact structure of NCP compared to DNA.

**Table 2.**
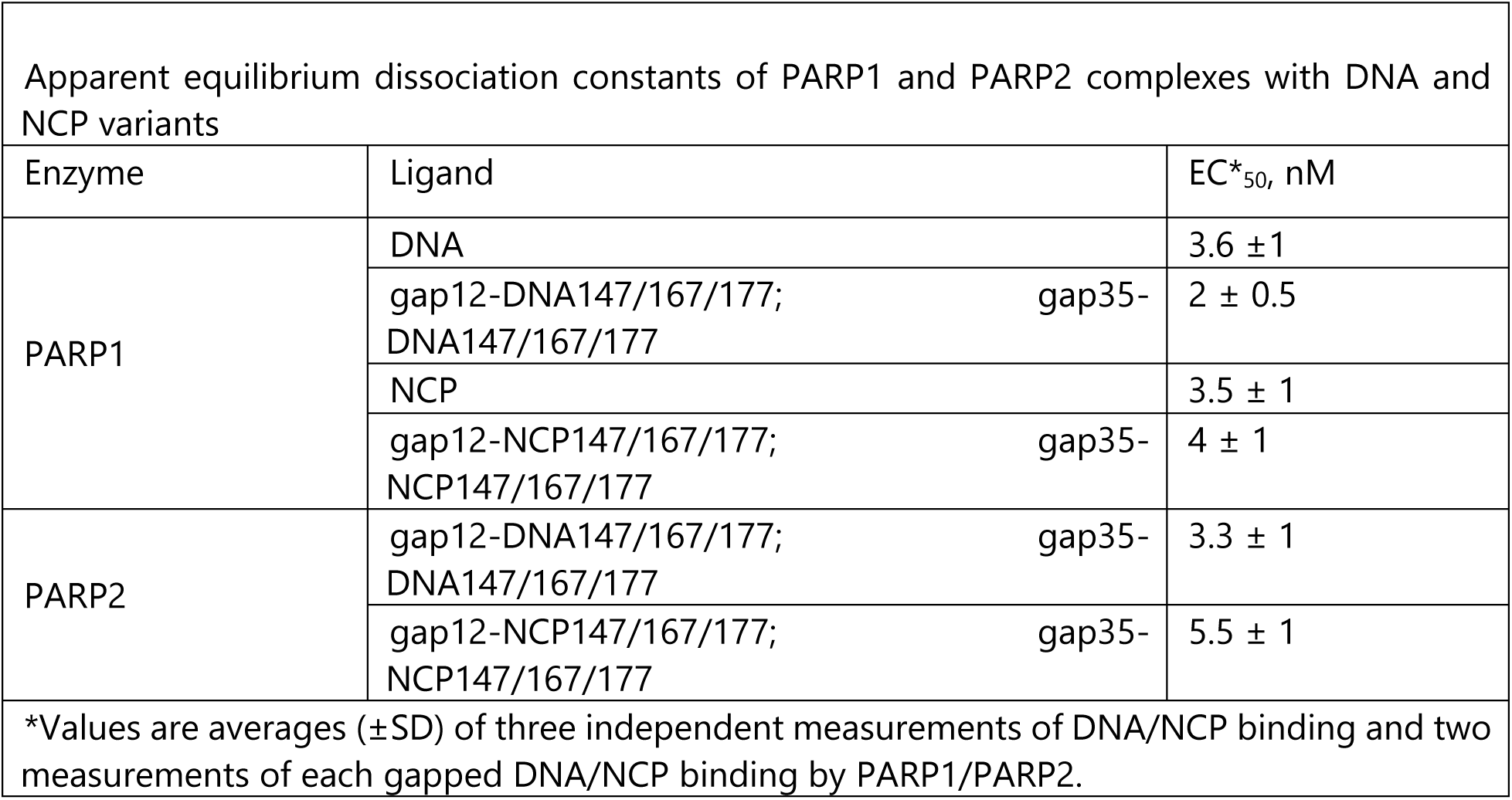
Apparent equilibrium dissociation constants of PARP1 and PARP2 complexes with DNA and NCP

Binding of PARP2 to nongapped DNA/NCP at the protein concentrations used in these experiments could not be quantified due to the absence of saturation (Supplementary Fig. 1). Similar to PARP1, PARP2 displayed practically identical affinities for all structural variants of gap-NCP, which were about 1.5-fold lower (*p*˂0.002) than respective values for PARP1. It should be noted that the lowest EC_50_ value (2 nM) is near the minimal concentration of fluorescently labelled ligand we used to perform reliable measurements and therefore represents the upper limit. It is very likely that this method limitation results in the absence of significant difference between the gapped DNA variants in the EC_50_ values.

The inability to distinguish the affinity of PARP1 for different gap-DNAs in direct titration experiments prompted us to perform further experiments using a competition binding assay. Plasmid DNA containing no breaks was added at increasing concentrations as the undamaged competitor to the preformed PARP1 complex with a defined DNA structural variant. Release of the fluorescently labelled DNA from the complex due to binding of the competitor is accompanied by a decrease in fluorescence anisotropy (Fig. 4a), enabling to calculate CC_50_ value (directly proportional to stability of the preformed complex). These experiments revealed that the strength of PARP1-DNA interaction depends on the presence of gap and its position relative to the blunt ends of all length variants (Fig. 4b).

**Fig. 4.**
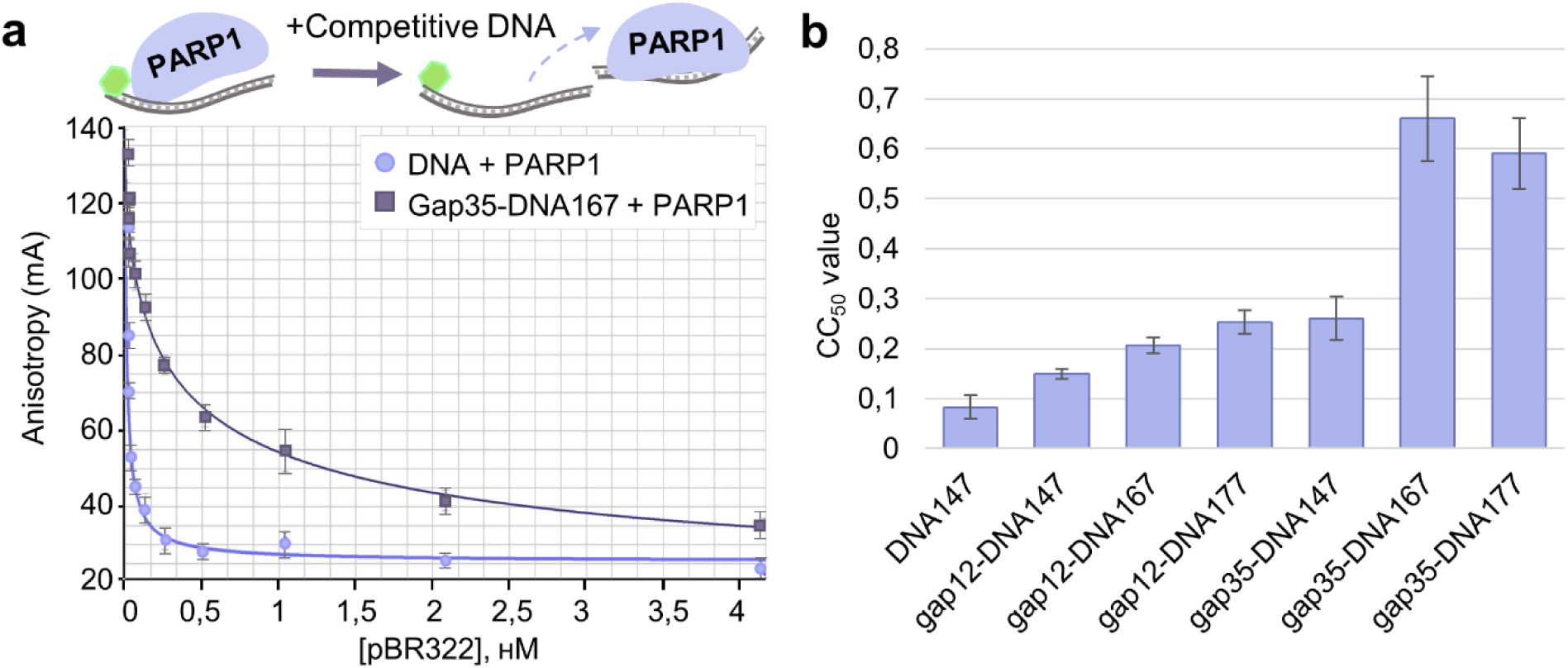
PARP1 affinity for gap-containing DNA depends of the damage position relative to the blunt ends: **a** − Typical titration curves reflecting dissociation of fluorescently labelled DNA147 or gap35-DNA167 from the complex with PARP1 upon addition of increasing concentrations of the competitor DNA (plasmid pBR322); **b** − CC_50_ values determined for different DNA structural variants.

Using direct fluorescence titration experiments, we revealed only a slight difference between PARP1 and PARP2 in the affinities for gapped NCPs (Table 2). However, their complexes can differ in composition due to propensity of both enzymes for self-association. To test this assumption, we performed mass photometry (MP) experiments. This novel technique measures the mass of individual molecules in solution and allows to obtain molecular mass distribution of proteins and other biomolecules reflecting the molecular composition of the sample and to measure binding affinity. MP experiments with individual components revealed coexistence of monomeric and dimeric forms of PARP1/PARP2 and coexistence of NCP147 with free DNA147 (Fig. 5a). PARP2 detected as more prone to dimerization compared to PARP1 was capable even of trimerization.

**Fig. 5.**
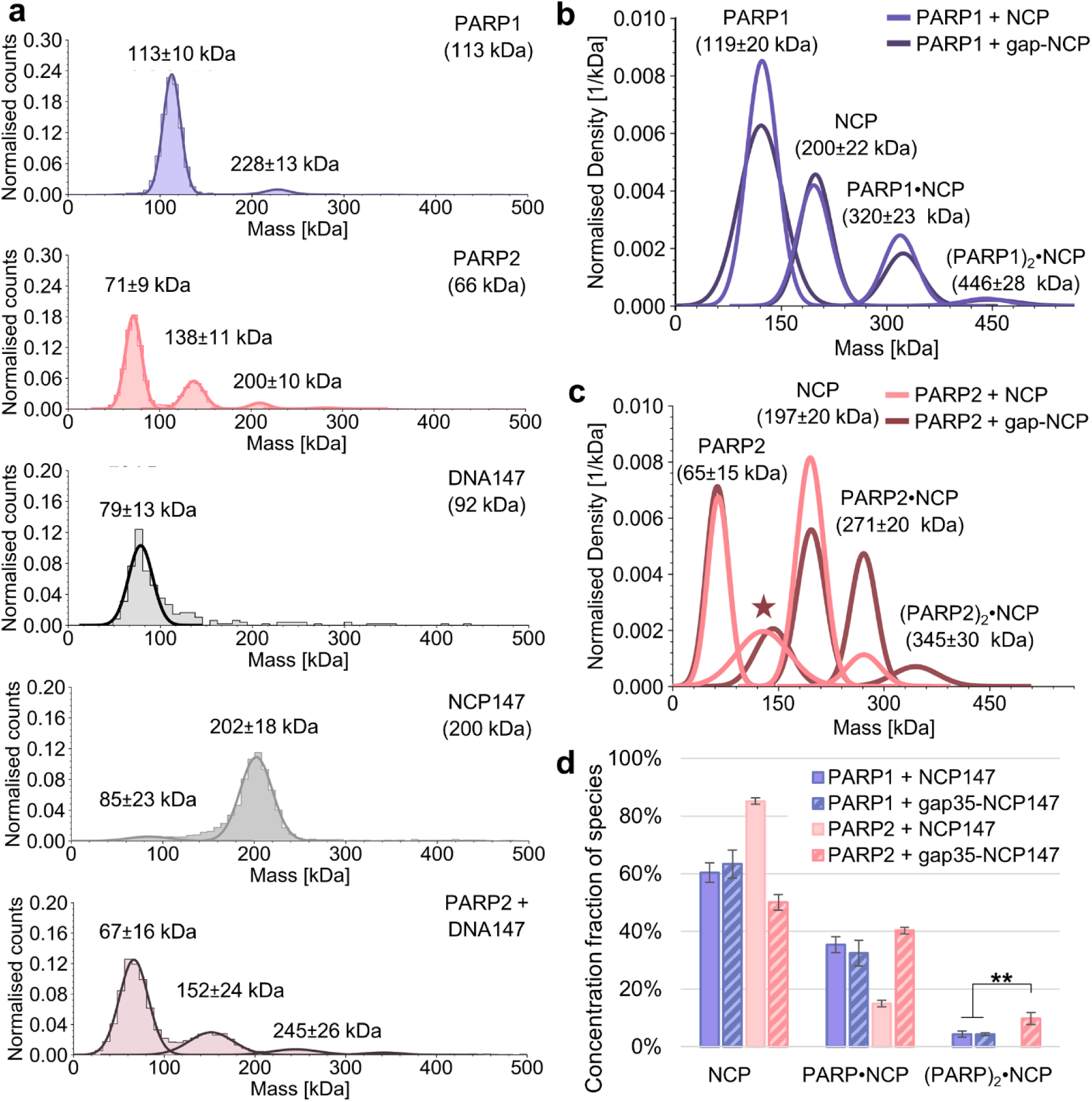
Mass photometry data: **a** – control measurements of samples containing individual components specified in each panel, **b, c** – molecular mass distribution of species in PARP1/PARP2 (6 nM) mixtures with NCP147/gap35-NCP147 (3 nM), **d** – histograms show relative amounts of species in the mixtures; difference between the relative amounts of (PARP1)_2_•gap-NCP and (PARP2)_2_•gap-NCP complexes was statistically significant (\*\**p* < 0.01).

PARP1 was revealed to form complexes with NCP147/gap35-NCP147 containing one or two protein molecules, PARP1•NCP (320±23 kDA) and (PARP1)_2_•NCP (446±28 kDa), respectively (Fig. 5b). In addition to analogous complexes formed by PARP2, PARP2•NCP (271±20 kDa) and (PARP2)_2_•NCP (345±30 kDa), a 130−142 kDa species was detected, which could correspond to PARP2 dimer or PARP2 complex with free DNA (Fig. 5c, the asterisked peak, Supplementary Fig. 2).

From the data of MP experiments, K_D_ values of PARP1/PARP2 complexes with NCP147 and gap35-NCP147 were determined, using method described by Wu and Piszczek (46). These values were calculated for the two different types of complexes formed by PARP1/PARP2 (Table 3), based on relative amounts of NCP in free and protein-bound states. Their comparison shows that the non-gapped NCP is more tightly bound by PARP1 than by PARP2. PARP2 has a higher affinity for gap-NCP: K_D_ values of PARP2•gap-NCP and (PARP2)_2_•gap-NCP complexes are 2−2.5-fold lower compared to the respective values of PARP1 complexes. On the contrary, PARP1 revealed a higher affinity for gap-NCPs in fluorescence titration experiments (Table 2).

**Table 3.** K_D_ values of PARP1 and PARP2 complexes with NCP147 and gap35-NCP147 (nM)

| Enzyme•NCP | Type of complex |  |
| --- | --- | --- |
|  | PARP•NCP | (PARP)<sub>2</sub>•NCP |
| PARP1•NCP147 | 8 ± 1 | 40 ± 10 |
| PARP1•gap35-NCP147 | 9 ± 2** | 40 ± 5*** |
| PARP2•NCP147 | 33 ± 4 | - |
| PARP2•gap35-NCP147 | 5 ± 1** | 16 ± 3*** |
| The K<sub>D</sub> values are averages (±SD) of three independent measurements. Statistically significant difference in K<sub>d</sub> values of PARP1 and PARP2 complexes with gap35-NCP147 is marked: $p < 0.01$(**) and $p < 0.001$ (***). | | |

The discrepancy between the data may result from the higher propensity of PARP2 for oligomerization. Binding of additional PARP2 molecules could contribute to an increase in the maximum fluorescence anisotropy level and revaluation of EC_50_ values. Furthermore, the PARP2 complex containing two protein molecules per NCP molecule was detected only for the gapped NCP, contrasting this enzyme to PARP1 (Fig. 5,c,d). Additional experiments with varied protein concentrations revealed dimerization of PARP2 in the complex with nongapped NCP at significantly higher protein concentration, suggesting dependence of the enzyme oligomerization capacity on the mode of interaction with NCP (Supplementary Fig. 3).

## DISCUSSION

### The Architecture of Catalytically Active PARP1/PARP2 Complex with NCP Impacts Histone Modification

Our results demonstrate that HPF1-dependent ADP-ribosylation of histones is influenced by the position of base excision repair (BER) specific DNA damage within the nucleosome core particle. Both PARP1 and PARP2 catalyze the modification of histones in NCP structures containing the one-nucleotide gap at position 35 more efficiently compared to position 12, regardless of the linker length (gap35-NCP147/167/177 vs gap12-NCP147/167/177, Fig. 1 and 3). We observed a ∼1.5-fold increased level of histones modification in gap35-NCPs despite the absence of detectable difference in the enzymes’ affinities for gap35-NCPs vs gap12-NCPs (Table 2). This suggests that mutual location of the enzyme binding site in DNA and ADP-ribose acceptors in core histones is critically important for the heteromodification reaction. On the other hand, the balance between auto- and heteromodification reactions catalysed by PARP2 strongly depends on the presence of gap in nucleosomal DNA (as evidenced from comparison of these data for NCP and gap12-NCPs or gap35-NCPs, Fig. 3,c), which is necessary for the high-affinity enzyme interaction with NCP. Thus, both the affinity of PARP for NCP and the architecture of catalytically active PARP-NCP complex determine the efficiency of histones modification. This conclusion is further supported by experiments with PARP2 and gapNCP267 bearing gap in the linker region: a significant reduction in the histone modification level (compared to that in the presence of NCP147/NCP267, Fig. 1,d), can be caused by PARP2 binding far away from the nucleosome core. We have also shown that preferable targets of PARP2 catalytic activity are histone tails located in close proximity to the DNA gap (Fig. 2), which may be crucial in generating a local DNA damage signal and preventing excessive chromatin ADP-ribosylation. Therefore, we propose that the DNA gap in the NCP serves as the main binding site for PARP2, coinciding with our previous data (13,32), and the enzyme modifies targets in close proximity to this binding and activation site on DNA.

Heteromodification of histones catalysed by PARP1 was also revealed to depend to different extents on the DNA gap presence and its position (Fig. 1, c), while no difference in the EC_50_ values of PARP1 complexes with different gapped NCPs was detected (Table 2). At the same time, the affinity of PARP1 for free gapped DNA duplexes appeared to depend on the distance between the gap and the blunt ends (Fig. 4,b) as potential binding sites. It seems reasonable to hypothesize existence of a specific binding site for PARP1 in the NCP. Previous *in vitro* studies have shown that PARP1 competes with linker histone H1 for binding near the entry-exit site of NCP, interacts with H3 and H4 histones and is capable of remodelling the NCP structure upon the binding (52−56). Furthermore, very early studies indicated predominant localization of PARP1 within internucleosomal linker regions of HeLa cell chromatin [for review see (18)]. This specific mode of PARP1 binding to NCP visualized recently by atomic force microscopy (57) can explain the effects we observed. Minor differences in PARP1-catalysed histone modification levels were noted for all NCPs utilized, excepting gap35-NCPs (discussed later). Gap introduction into the linker region did not result in a significant decrease in histone modification by PARP1, and localization of the gap within the nucleosome core did not alter the histone modification pattern. Based on these findings, we suggest that PARP1 binds preferentially near the entry-exit region of NCPs, while the DNA gap acts as an alternative site for PARP1 binding responsible for the enhanced histone modification.

### ADP-ribosylation Targets in Nucleosome are Determined by Dynamics of Histones Surrounding the DNA Damage

PARP1 and PARP2 revealed dependence of the histone modification level and balance between the auto- and heteromodification reactions on the position of gap within the NCP structure (Fig. 3), despite of the difference between the enzymes in predominant modes of binding to NCP: near the entry-exit site (PARP1) or to the gap (PARP2). Both enzymes display very similar affinities for gap35-NCP and gap12-NCP (Table 2), indicating independence of the extent of histone modification on the strength of PARP-NCP interaction. Furthermore, the superhelical location of the DNA damage determines the accessibility of PARP2 to the certain histones to be modified reflected in a specific pattern of heteromodification reaction (Fig. 2). We therefore suggest that the selectivity and efficiency of histone modification is primarily determined by the local environment of the DNA damage (acting as a PARP binding site) in the NCP structure and dynamics of histone-DNA interaction. Similar observations were published previously for DNA glycosylases. The enzymatic activities of UDG and OGG1 in NCP structures do not correlate with the solution accessibility of the DNA lesions but significantly depend on the NCP dynamics modulated by modification of histone tails (58,59). The molecular dynamics simulations study provides evidence that tails of core histones differ in the conformational flexibility and exhibit distinct binding modes to specific DNA regions (35). Interestingly, histone H3 characterized by the lowest flexibility of its arginine-rich tail is the worst ADP-ribosylation target of PARP1 and PARP2 (Fig. 2). The ability to modify histone H3 detected solely for PARP1 can be explained by the fact that predicted from simulations multiple DNA-interaction sites for H3 tails are around the primary binding site of the enzyme (the entry-exit site). Our data allow to suggest that BER-specific DNA lesions localized near SHL 3.5 of NCP may represent hot points to promote ADP-ribosylation-dependent chromatin relaxation.

### Mass Photometry Allows to Analyze the Stoichiometry of PARP-NCP Complexes and Self-association of PARPs

Mass photometry (MP), a relatively novel technique, enables to detect and accurately measure molecular masses of proteins, nucleic acids, and their complexes (46). Here, we applied the MP measurements to analyze formation and composition of PARP-NCP complexes and the oligomeric state of PARPs in solution. Considering the self-association of PARP1/PARP2 and their affinity for undamaged DNA, we performed MP experiments at PARP/NCP concentrations close to EC_50_ values of their complexes, as determined by the fluorescence anisotropy measurements. Our results revealed the presence of species corresponding to (PARP)_2_•NCP complexes even at low concentrations with only a two-fold excess of PARP over NCP (6 nM and 3 nM, respectively). The patterns of species distribution observed for PARP1 mixtures with NCP147 and gap35-NCP147 were very similar, indicating low contribution of PARP interaction with gap to the complex formation as suggested from ADP-ribosylation experiments. Both MP and fluorescence anisotropy measurements of binding affinities revealed distinction between PARP1 and PARP2: the strength of interaction with NCP strongly depends on the presence of gap only for PARP2 (Tables 2 and 3). Another difference evidenced from the MP experiments is significantly higher stability of PARP2 complex with gap35-NCP containing two protein molecules (divalent binding) compared to the respective complex of PARP1 (Table 3).

Self-association of PARP1 and PARP2 have been detected by different approaches in previous studies (60−63). Our MP experiments provide the first evidence of a higher propensity of PARP2 for oligomerization via di- and trimerization (Fig. 5a). More importantly, binding of two PARP2 molecules to one NCP molecule has never been detected. The cryo-electron-microscopic structural study of PARP2-HPF1 complex bound to mononucleosome revealed bridging of two NCPs via interaction of one or two PARP2 molecules with the ends of both DNAs (64). Cooperative binding of PARP1 to different structures of free DNA is well known (65−67). Binding of two or three PARP1 molecules to NCP depends on the presence of linker DNA (56). The atomic force microscopy study enabled to visualize dimerization of PARP1 and PARP2 at distinct damage types of free DNA, with dimer formation upon the interaction with gap being unique for PARP2 (68). All the data combined allow to suggest that the (PARP2)_2_•gap-NCP complex detected is composed of PARP2 dimer bound to the DNA gap.

## CONCLUSION

The details of PARP1 and PARP2 interaction with free DNA and nucleosome have been the subject of numerous studies, with some of them being focused on the relationship between the DNA-dependent enzymatic activity and affinity for the distinct DNA structure. The interrelation between the nucleosome (NCP) architecture and PARP-catalysed modification of histones as the earliest targets of ADP-ribosylation in response to DNA damage has never been studied by others. Here, by using a combination of biophysical and biochemical approaches we succeeded to reveal that the extent of histone ADP-ribosylation and the balance between auto- and heteroPARylation reactions depend to different extents for PARP1 and PARP2 on the presence of a one-nucleotide gap (the BER-specific lesion) and its location in the NCP structure (Fig. 6). A higher dependence of the level and pattern of PARP2-catalysed histone modification on the NCP architecture is easily explained by the gap-binding induced enzyme activation. This specific PARP2-NCP interaction favors the enzyme functioning as a dimer. No detectable impact of the NCP structure on selectivity of histone modification in the case of PARP1 obviously results from the predominant enzyme binding at the entry-exit region. The analysis of histone modification data taking into account the known HPF1-dependent ADP-ribosylome of histones and dynamics of the NCP structure revealed interrelation between the pattern and extent of histone modification and conformational flexibility of the target histone tails. The location of a specific DNA damage and dynamics of NCP modulated by various post-translational modifications may determine distinct roles of PARP1 and PARP2 in response to genotoxic stress.

**Fig. 6.**
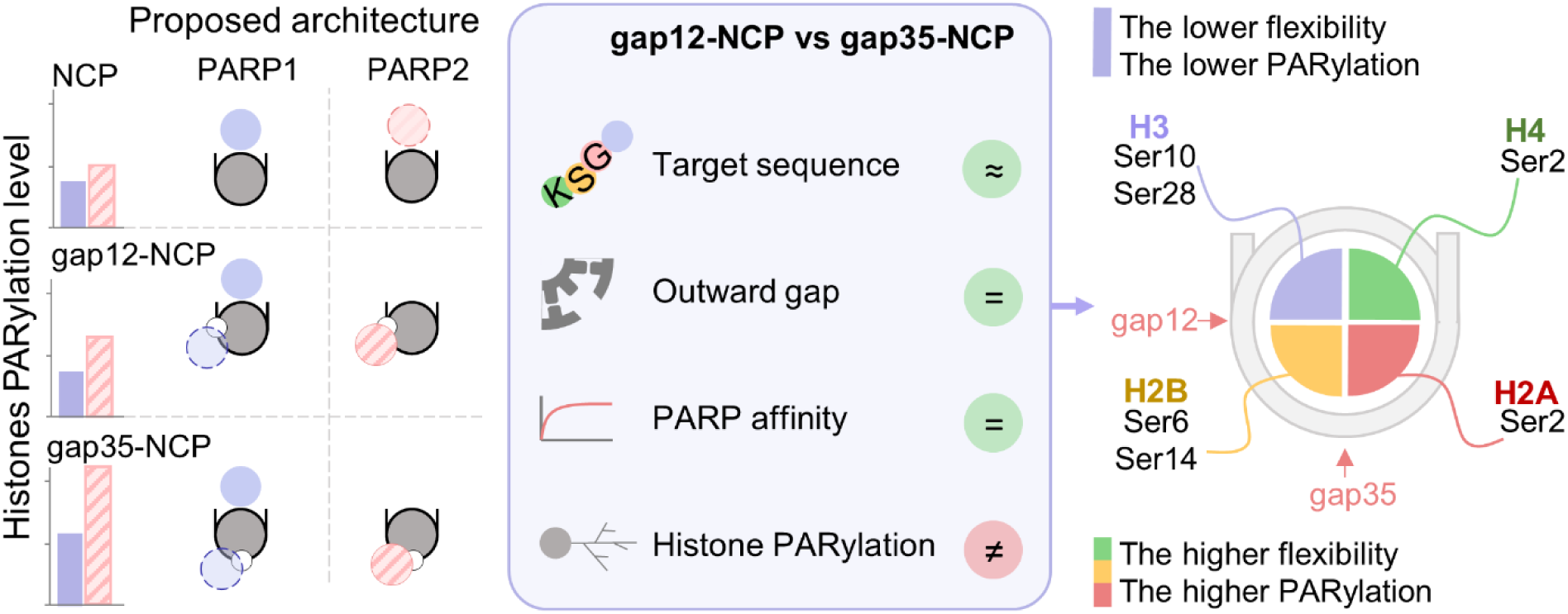
Histone PARylation in nucleosome is controlled by the specificity of PARP-NCP interaction and conformational flexibility of histone tails. The presence and position of the one-nucleotide gap in nucleosomal DNA were revealed to produce a significant effect on the extent of histone PARylation despite of the absence of detectable difference in the PARP1/PARP2 affinities for the two gapped NCP structures. We suggest that superhelical location of the BER-specific DNA lesion as a potential binding site for PARP activation and the conformational dynamics of surrounding histone tails determine the efficiency of histone ADP-ribosylation and following relaxation of NCP structure. The known sites of serine ADP-ribosylation on histones are specified in accordance with (25).

## AUTHORS’ CONTRIBUTIONS

K.T.A. investigation; M.N.A., and K.T.A. conceptualization, methodology and writing; K.M.M. resources; E.A.V., and K.T.A. data curation and validation; L.O.I. conceptualization, writing and supervision.

## ACKNOWLEDGEMENTS

We would like to thank the entire laboratory of bioorganic chemistry of enzymes for feedback. We acknowledge Konstantin N. Naumenko and Alexander A. Ukraintsev for preparation of recombinant HPF1, and Dmitry O. Zharkov for supporting Mass Photometry experiments.

The online version contains supplementary material available at The data underlying this article will be shared on reasonable request to the corresponding author.

