## Supplementary material for "DECIPHERING THE DARK SIDE OF HISTONE ADP-RIBOSYLATION: WHAT STRUCTURAL FEATURES OF DAMAGED NUCLEOSOME REGULATE THE ACTIVITIES OF PARP1 AND PARP2"

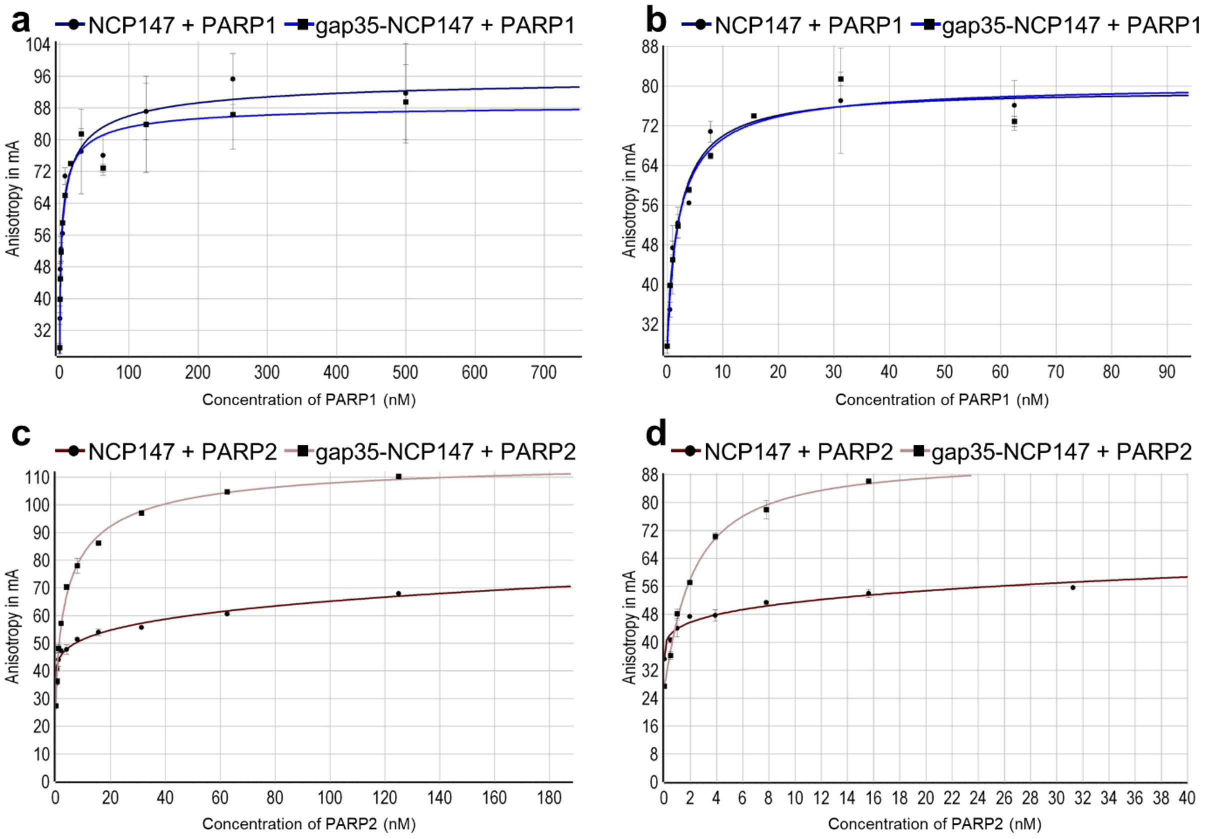


**Supplementary figure 1.** Typical titration curves representing binding of PARP1 (**a,b**) and PARP2 (**c,d**) to NCP147/35gap-NCP147 (3 nM) at different protein concentration range, obtained by measurements of FAM fluorescence polarization.

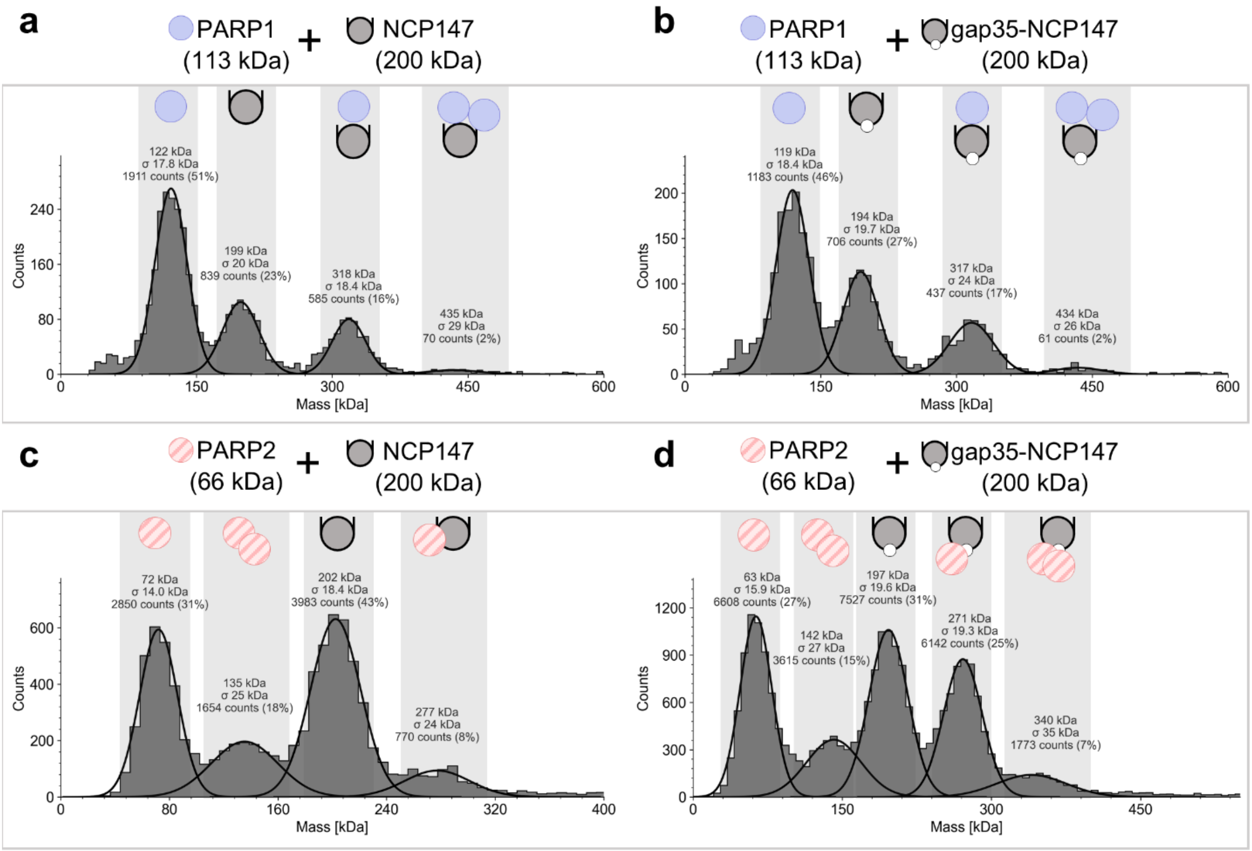


**Supplementary figure 2.** Representative mass photometry data reflecting complex formation upon binding of PARP1 (6 nM) (**a**,**b**) and PARP2 (6 nM) (**c, d**) to NCP147/gap35-NCP147 (3 nM). The species detected are schematically shown in the panels.

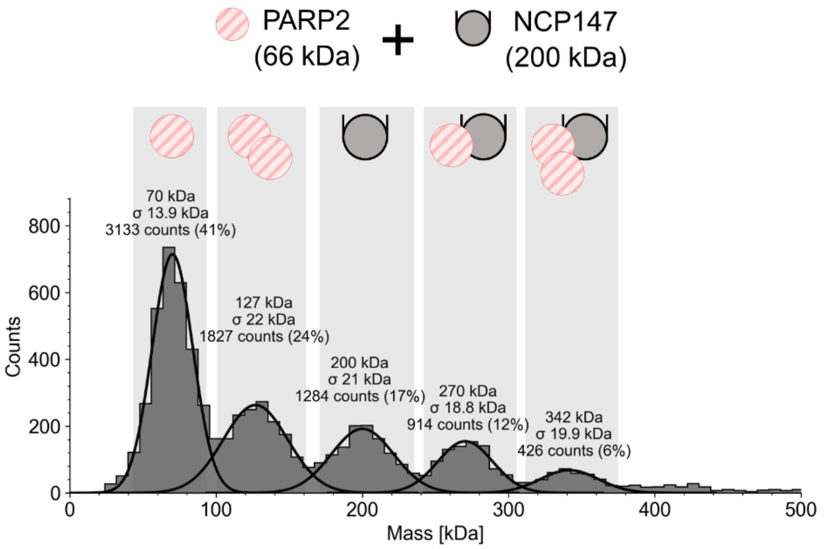


**Supplementary figure 3.** Mass photometry analysis of PARP2 dimerization in the complex with nongapped NCP. Representative mass photometry data reflecting distribution of species formed in the mixture of PARP2 (20 nM) with NCP147 (5 nM).
